# A multivalent docking platform and Rcn1-mediated inhibition control the extent of calcineurin recruitment to the cell division site for the dephosphorylation of multiple cytokinetic proteins

**DOI:** 10.64898/2026.09.14.751435

**Authors:** Alaina H. Willet, Jun-Song Chen, Qing Yu, Chloe E. Snider, Liping Ren, Rahul Bhattacharjee, Steven P Gygi, Kathleen L. Gould

**Affiliations:** Department of Cell and Developmental Biology, Vanderbilt University School of Medicine, Nashville, TN 37232, USA; Department of Cell Biology, Harvard Medical School, 240 Longwood Ave, Boston, MA 02115, USA; Department of Biochemistry and Molecular Biotechnology, University of Massachusetts Chan Medical School, Worcester, MA 01605

## Abstract

Cytokinesis requires coordinated signaling to ensure the accurate physical separation of daughter cells. Calcineurin (CN), a conserved Ca²⁺/calmodulin-dependent phosphatase, is required for cytokinesis in organisms ranging from yeast to humans, yet how CN is regulated at the division site and the substrates through which it promotes cell division remain poorly understood. Here we use the fission yeast *Schizosaccharomyces pombe*, which display striking cell division defects in the absence of CN, to define how CN is anchored at the cell division site. We show that CN recruitment to the cytokinetic ring (CR) requires its PxIxIT- and LxVP-binding surfaces and is mediated by multivalent interactions with the CR components paxillin-like Pxl1 and the F-BAR protein Cdc15. Disrupting these interactions nearly eliminates CN from the CR and causes gross cytokinetic defects similar to complete loss of CN function. Cell cycle stage-specific quantitative phosphoproteomics combined with proximity labeling-based proteomics were used to identify candidate CN substrates involved in cytokinesis. Validation of a cohort of these proteins localizing to the CR, including the F-BAR protein Rga7, the actin regulator Aim21 and three protein kinases, revealed that CN targets a broad network of structural and signaling components involved in cell division. We also identify the conserved CN inhibitor Rcn1 as a CN substrate and show that Rcn1 restricts CN accumulation at the CR to provide an additional layer of spatial regulation. Thus, spatial control of CN enables proper protein dephosphorylation for successful cytokinesis.

## Introduction

Cytokinesis is the final stage in the cell cycle that results in the production of two new daughter cells. In many eukaryotes, this process is driven by the assembly and constriction of an actomyosin-based cytokinetic ring (CR) that is anchored at the plasma membrane and constricts to result in daughter cell separation (Glotzer, 2017). The fission yeast *Schizosaccharomyces pombe* has been a powerful model for uncovering conserved mechanisms of cytokinesis due to its symmetrical division, tractable genetics, and reduced levels of redundancy (Vyas et al., 2021). Large-scale and targeted studies have cataloged approximately 50 proteins that localize to the CR in *S. pombe* (Carme et al., 2026). Despite this comprehensive inventory of cytokinetic components, the signaling mechanisms that regulate their activity and coordination remain incompletely understood.

One critical signaling input during *S. pombe* cytokinesis is the conserved serine/threonine phosphatase calcineurin (CN), a Ca²⁺/calmodulin-dependent enzyme (Yoshida et al., 1994). CN localizes to the CR, and *S. pombe* cells lacking CN function exhibit delayed CR constriction accompanied by thickened septa, resulting in chains of branched, incompletely separated slow-growing multi-septated cells (Yoshida et al., 1994; Lu et al., 2002; Kozubowski et al., 2011b; Martín-García et al., 2018; Snider et al., 2020). In human cells, CN also localizes to the midbody, the site of abscission, and loss of CN function leads to multinucleation due to failed cytokinesis (Chircop et al., 2010). However, the mechanism by which CN promotes the final step of cell division is not well understood in any organism.

Calcineurin is composed of a catalytic subunit and a regulatory subunit, and this complex can also associate with calmodulin (Li et al., 2011; Ulengin-Talkish and Cyert, 2023). The catalytic subunit is kept in an autoinhibited state via a C-terminal autoinhibitory sequence (AIS), which is released upon Ca²⁺-calmodulin binding (Hashimoto et al., 1990; Stemmer and Klee, 1994; Kissinger et al., 1995; Perrino et al., 1995; Li et al., 2016; Chen, 2025). In contrast to humans and *Saccharomyces cerevisiae*, in which multiple genes encode each CN subunit, *S. pombe* contains a single gene for each, simplifying genetic dissection of CN function (Cyert et al., 1991; Yoshida et al., 1994; Ulengin-Talkish and Cyert, 2023; Rutherford et al., 2024). This makes *S. pombe* a particularly advantageous system for studying CN signaling during cytokinesis.

CN recognizes substrates through short linear motifs (SLiMs), including the PxIxIT and LxVP motifs, which dock onto specific regions of the enzyme (Roy and Cyert, 2009; Grigoriu et al., 2013; Nygren and Scott, 2016). In mammals, several well-characterized CN substrates harbor these motifs, including NFAT transcription factors and the NOTCH1 receptor (Matheos et al., 1997; Stathopoulos and Cyert, 1997; Hirayama et al., 2003; Ly and Cyert, 2017; Wigington et al., 2020). Dynamin is the only known CN substrate at the division site in human cells, and no additional cytokinesis-specific substrates have been identified (Chircop et al., 2010).

In *S. pombe*, the F-BAR protein Cdc15 is currently the only confirmed CR-localized CN substrate (Martín-García et al., 2018; Snider et al., 2020). Cdc15, comprising an N-terminal F-BAR domain, a central intrinsically disordered region (IDR), and a C-terminal SH3 domain, functions as a major scaffold for the CR (reviewed in (Snider et al., 2021)). It contains more than 30 phosphorylation sites, primarily clustered in the IDR (Roberts-Galbraith et al., 2010; Swaffer et al., 2016; Lee et al., 2018; Bhattacharjee et al., 2020, 2023; Magliozzi et al., 2020). These sites are phosphorylated during interphase by multiple kinases and undergo dephosphorylation at mitotic entry, a transition required for Cdc15 membrane condensation, oligomerization, protein partner binding, and CR assembly (Wachtler et al., 2006; Roberts-Galbraith et al., 2009, 2010; Arasada and Pollard, 2014; Cortés et al., 2015; McDonald et al., 2015; Ren et al., 2015; Willet et al., 2015; Snider et al., 2020, 2022; Bhattacharjee et al., 2023). Mutations that mimic or block phosphorylation impair Cdc15 function and exacerbate defects in other cytokinesis mutants, underscoring the importance of its phosphoregulation (Roberts-Galbraith et al., 2010; Bhattacharjee et al., 2020, 2023).

Beyond serving as a substrate, Cdc15 contributes to CN localization at the CR. A Cdc15 mutant lacking part of the IDR (Cdc15-Δ2) displays a strong reduction in CN localization at the division site (Mangione et al., 2019). The paxillin-like protein Pxl1 is also implicated in CN recruitment, though it is not a CN substrate and remains correctly localized in *cdc15-Δ2* cells (Martín-García et al., 2018; Mangione et al., 2019). These observations led to a model in which Cdc15 and Pxl1 cooperate to tether CN to the CR, enabling efficient substrate dephosphorylation (Martín-García et al., 2018; Snider et al., 2020). However, the mechanism of this cooperation was unknown.

In this study, we determine the molecular mechanisms underlying CN localization and function at the CR. We show that substrate docking sites and catalytic activity are essential for CN accumulation at the division site and for its role in cytokinesis. We define how Cdc15 and Pxl1 collaborate to assemble a multivalent platform for CN recruitment. Because Cdc15 is not the sole CN substrate during cell division, cell cycle stage-specific quantitative phosphoproteomics and proximity labeling experiments were conducted and multiple CN substrates at the cell division site in *S. pombe* were validated. These findings expand our understanding of how CN coordinates cell division and uncover new regulatory substrates essential for cytokinesis.

## Results

### SLiM Docking Sites and Catalytic Activity Regulate Calcineurin Localization and Function at the Cytokinetic Ring

To assess the contribution of SLiM docking sites to CN function during *S. pombe* cytokinesis, we generated point mutations in the CN catalytic subunit, Ppb1, that disrupt known substrate interaction motifs (Li et al., 2004; Roy et al., 2007; Rodríguez et al., 2009; Grigoriu et al., 2013). The PxIxIT docking site was mutated (N359A, I360A, R361A; *ppb1-3A*), the LxVP docking site was mutated (W371A; *ppb1-W371A*), and a combined mutant targeting both sites was also constructed (*ppb1-4A*) (Fig 1A). Each allele was tagged with the sequences encoding mNeonGreen (mNG) and compared to wildtype CR localization. Ppb1-mNG lacking either the PxIxIT or LxVP docking site had reduced CR localization compared to wildtype (Fig 1B and 1C) and Ppb1-4A-mNG was absent from the cell division site (Fig 1B and 1C). To determine whether docking site disruption impaired CN function, we quantified septation defects, a hallmark of CN loss-of-function (Yoshida et al., 1994). Wildtype and *ppb1-W371A* showed normal cell phenotypes while *ppb1-3A* showed a modest increase (∼3%) in cells with two or more septa (Fig 1D and 1E). In contrast, *ppb1-4A* phenocopied *ppb1Δ*, with ∼10% of cells showing cell separation defects (Fig 1D and 1E). These results indicate that SLiM docking sites are critical for CN localization to the CR and are required for its function in cytokinesis.

**Fig 1.**
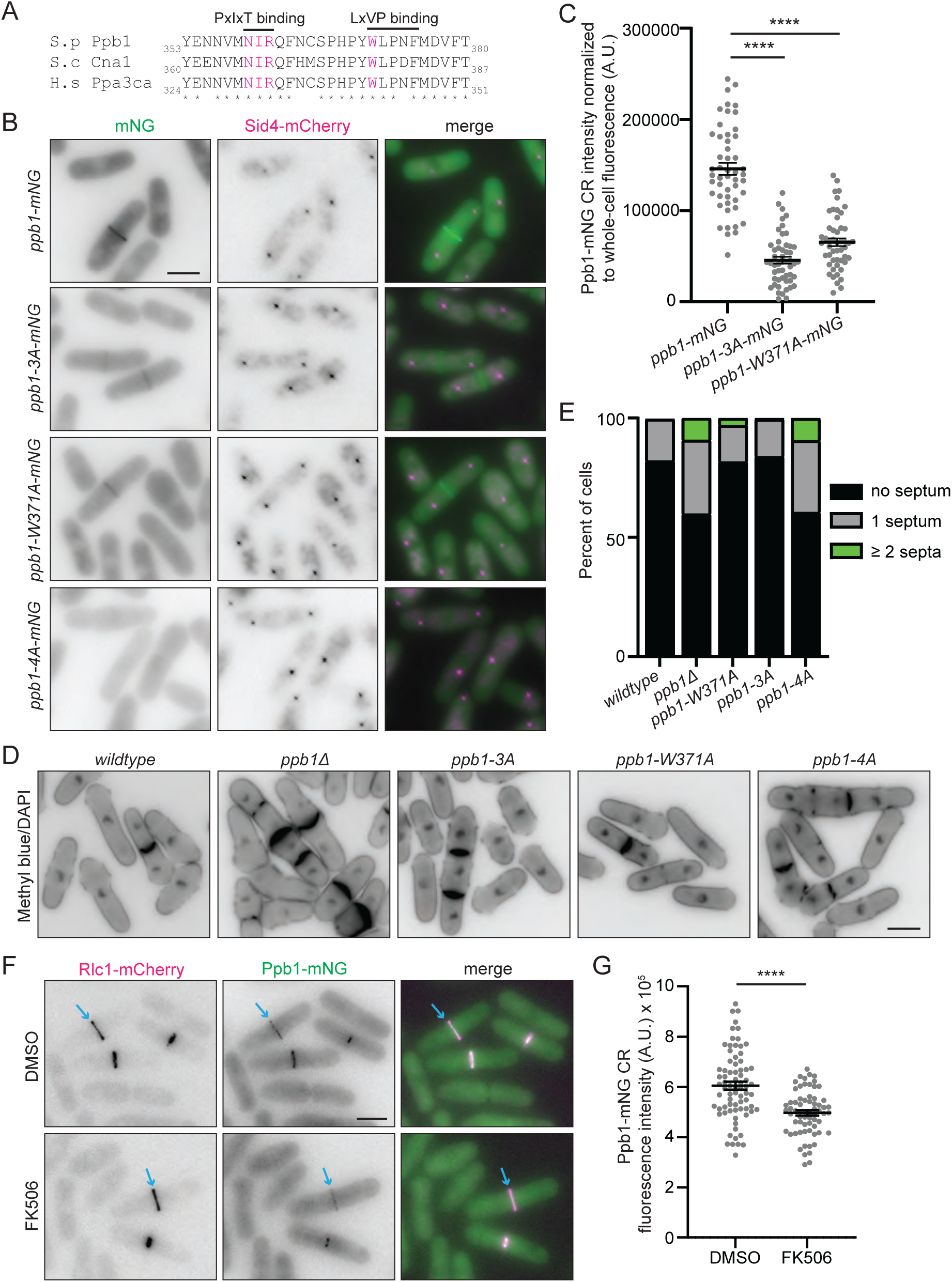
Calcineurin substrate docking sites are required for CR localization and function. **(A)** Sequence alignment highlighting residues involved in PxIxIT and LxVP binding; conserved residues are marked with asterisks. The residues mutated in the docking mutants are highlighted in magenta. **(B)** Live-cell images of cells expressing wildtype or mutant Ppb1-mNeonGreen (mNG) proteins. Sid4-mCherry marks spindle pole bodies and mitotic cells. **(C)** Quantification of Ppb1-mNG fluorescence at the CR, normalized to total cellular fluorescence. *n* ≥ 50 cells from two biological replicates. \*\*\*\**p* < 0.0001 by unpaired two-tailed Student’s *t*-test. **(D)** Fixed-cell images of the indicated strains stained with DAPI and Methyl Blue following growth at 25°C in YE. **(E)** Quantification of septation phenotypes from D. *n* ≥ 578 for each strain from two biological replicates. **(F)** Live-cell imaging of Ppb1-mNG Rlc1-mCherry cells treated for 10 minutes with 10 μg/ml of FK506 or DMSO prior to imaging. Blue arrows indicate examples of fully formed CRs that have not begun constriction. **(G)** Quantification from F of Ppb1-mNG intensity at the CR, normalized to Rlc1-mCherry signal. *n* ≥ 38 cells from three biological replicates. \*\*\*\**p* < 0.0001 by unpaired two-tailed Student’s *t*-test. Scale bars = 5 μm.

To further probe the importance of LxVP docking, we treated cells with the CN inhibitor FK506, which blocks LxVP-mediated substrate binding (Grigoriu et al., 2013). Short-term treatment (10 minutes) reduced Ppb1-mNG CR localization by ∼50%, without affecting the CR marker Rlc1-mCherry (Le Goff et al., 2000; Naqvi et al., 2000). (Fig 1F and 1G). These results are consistent with prior long-term inhibitor treatments (Martín-García et al., 2018) and support a model in which both PxIxIT and LxVP docking interactions contribute to stable CN recruitment to the CR.

We next asked if CN’s phosphatase activity influences its localization. For this, we generated two activity-altering *ppb1* alleles based on human disease-associated mutations: *ppb1-N179I*, which impairs phosphatase activity, and *ppb1-F499L*, which disrupts autoinhibition mediated by the AIS and renders CN hyperactive (Mizuguchi et al., 2018). Both Ppb1 mutant proteins were produced at similar levels to wildtype, and the cells showed the expected change in localization of transcription factor Prz1, an established CN substrate that translocates into the nucleus upon CN-dependent dephosphorylation (Hirayama et al., 2003; Mizuguchi et al., 2018). As expected, *ppb1-N179I* cells showed septation defects similar to *ppb1Δ*, while *ppb1-F499L* cells resembled wildtype (Fig 2A and 2B). We next tagged Ppb1-N179I and Ppb1-F499L with mNG in cells that also expressed Rlc1-mCherry and compared their localization to wildtype. Ppb1-N179I-mNG displayed a ∼45% reduction in CR localization, while Ppb1-F499L-mNG showed a ∼33% increase in CR localization compared to wildtype (Fig 2C and 2D).

**Fig 2.**
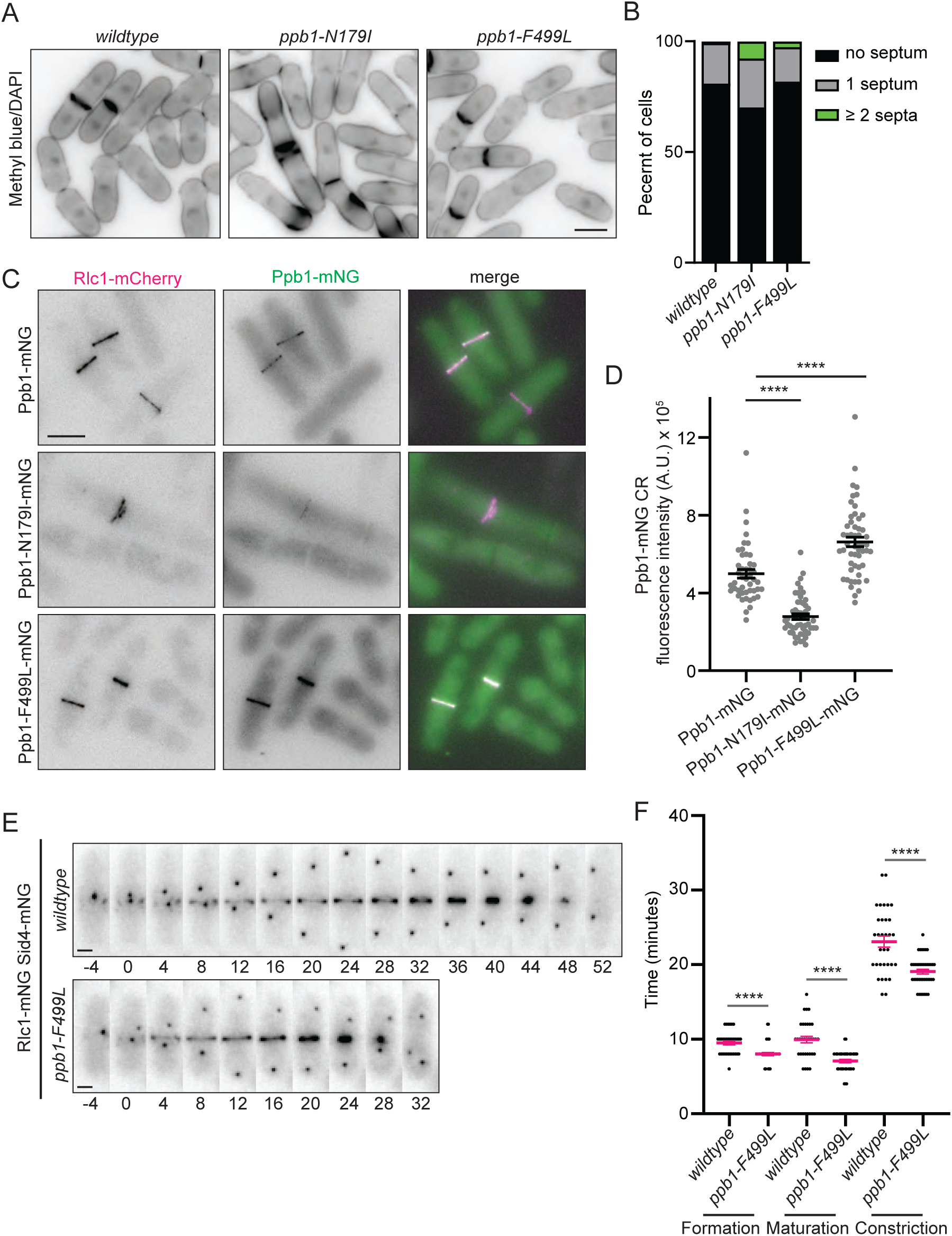
Calcineurin phosphatase activity is required for robust localization to the CR. **(A)** Fixed-cell images of the indicated strains grown at 25°C in YE and stained with DAPI and Methyl Blue. Scale bar = 5 μm. **(B)** Quantification of the septation phenotypes from A. *n* ≥ 338 for each strain. **(C)** Live-cell images of cells expressing Ppb1-mNG, Ppb1-N179I-mNG or Ppb1-F499L-mNG with Rlc1-mCherry. Scale bar = 5 μm. **(D)** Quantification from C of Ppb1-mNG signal at the CR, normalized to the Rlc1-mCherry signal. *n* ≥ 46 from three biological replicates. \*\*\*\**p* ≥ 0.0001 by unpaired, two-tailed Student’s *t*-test. **(E)** Live-cell time-lapse images of wildtype or *ppb1-F499L* cells expressing Rlc1-mNG and Sid4-mNG. Images were acquired every 2 min and every 4 min is shown. Time on graphs is in minutes and time zero is the first frame of SPB septation. Scale bar = 2 μm. **(F)** Quantification of cytokinetic timing from time-lapse movies in E, including CR formation, maturation, and constriction. *n* = 33 for wildtype and *n* = 41 for the mutant. Magenta bars represent mean ± SEM. \*\*\*\**p* ≥ 0.0001 by unpaired, two-tailed Student’s *t*-test.

To test whether enhanced CN activity influences CR kinetics, we performed live-cell imaging of Rlc1-mNG and Sid4-mNG in wildtype and *ppb1-F499L* cells. Time-lapse imaging revealed that hyperactive CN accelerated multiple stages of cytokinesis, including CR formation, maturation, and constriction (Fig 2E and 2F). These findings indicate that CN activity positively regulates its own CR accumulation and promotes cytokinesis.

### The F-BAR Protein Cdc15 and Paxillin-like Pxl1 Cooperate to Recruit Calcineurin to the Cytokinetic Ring via Direct Interactions

CN fails to localize to the CR in *S. pombe* cells lacking either a segment of the intrinsically disordered region of Cdc15 (IDR2; amino acids 503–677) (Mangione et al., 2019) or CR-localized Pxl1 (Martín-García et al., 2018; Snider et al., 2020). These observations suggest that Cdc15 and Pxl1 collaborate to recruit CN to the division site, but the molecular basis of this mechanism has remained unclear.

We used AlphaFold3 (AF3) to model a potential interaction interface between CN and Cdc15 (Abramson et al., 2024). AF3 predicted that the LxVP docking pocket on the CN catalytic subunit Ppb1 binds a conserved LxVP-like motif within Cdc15 IDR2 (^650^LGAP^653^) (Fig 3A, 3B and 3C). Using recombinant GST-tagged Cdc15(441-927), which includes the LxVP-like motif, and recombinant CN (Ppb1-His₆, Cnb1, and Cam1) (Snider et al., 2020), we confirmed a direct interaction between Cdc15 and CN *in vitro* (Fig 3D).

**Fig 3.**
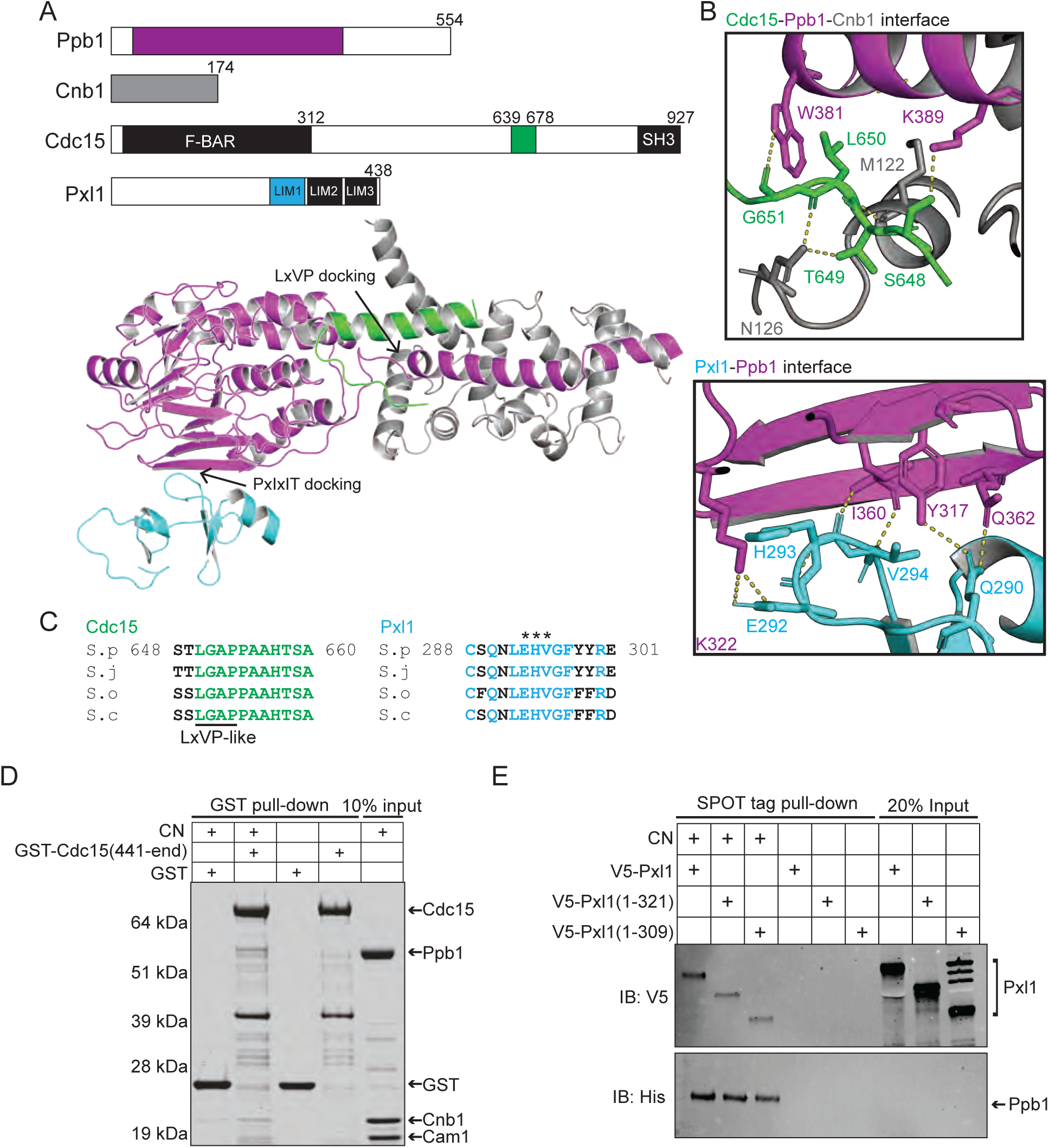
Cdc15 and Pxl1 directly bind calcineurin via distinct docking sites. **(A)** Scaled schematics of Ppb1, Cnb1, Cdc15, and Pxl1 (top), and AlphaFold3 structural models of the predicted Ppb1/Cnb1 complex bound to Cdc15 IDR2 (residues 648-678, green) and Pxl1 LIM1 (cyan). Ppb1 (residues 55-413) is shown in magenta, and Cnb1 in gray. **(B)** Close-up views of the predicted interaction interfaces between Ppb1 and Cdc15 (top), and Ppb1 and Pxl1 (bottom), from the model in A. **(C)** Sequence alignments of the Cdc15 and Pxl1 interaction regions across *Schizosaccharomyces* species: *S. pombe* (S.p), *S. japonicus* (S.j), *S. octosporus* (S.o), and *S. cryophilus* (S.c). **(D)** *In vitro* binding assay of recombinant CN (Ppb1-His₆, Cnb1, Cam1) incubated with bead-bound GST or GST-Cdc15(441-927). After 30 min incubation, samples were washed, resolved by SDS-PAGE, and stained with Coomassie. **(E)** *In vitro* binding assay of bead-bound CN (His₆-SPOT-Ppb1, Cnb1, Cam1) incubated for 1 h with V5-tagged full-length Pxl1, Pxl1(1-312), or Pxl1(1-309). After washing, samples were resolved by SDS-PAGE, transferred to membrane, and immunoblotted with anti-V5 and anti-His antibodies.

We also used AF3 to examine a potential Pxl1-CN interaction (Abramson et al., 2024). AF3 predicted that a conserved Pxl1 loop within the LIM1 domain bound the PxIxIT-binding surface on Ppb1 (Fig 3A and 3B). Notably, this Pxl1 loop lacks a canonical PxIxIT motif, suggesting an unconventional interaction. To test the AF3 prediction, we purified from bacteria full-length V5-tagged Pxl1 and two truncation mutants, V5-Pxl1(1-321) and V5-Pxl1(1-309), which lack LIM3 or both LIM2 and LIM3, respectively. All three proteins bound recombinant CN *in vitro* (Fig 3E), consistent with the LIM1 domain mediating the interaction, as predicted from the AF3 modeling.

To test the relevance of the above-described interactions *in vivo*, we constructed strains in which the ^50^LGAP^653^ motif in Cdc15 was mutated to AAAA (*cdc15-L650A,G651A,P653A*; *cdc15^LGAP*^*), or three residues in the Pxl1 LIM1 loop were mutated to alanine (*pxl1-E293A,H294A,V295A*; *pxl1^EHV*^*). We then analyzed Ppb1-mNG CR localization in these mutant backgrounds. Compared to wildtype cells, Ppb1-mNG levels were reduced by ∼33% in mCherry-c*dc15^LGAP*^* cells and ∼60% in *pxl1 ^EHV*^* cells compared to wildtype (Fig 4A and 4B). In the double mutant, CR-localized CN was reduced by ∼85% and was nearly undetectable at the CR in most cells (Fig 4A and 4B). Importantly, mCherry-Cdc15^LGAP*^ and mNG-Pxl1^EHV^* retained their CR localization, indicating that the loss of CN localization was not due to the absence of Cdc15 or Pxl1 at the CR (S1 Fig).

**Fig 4.**
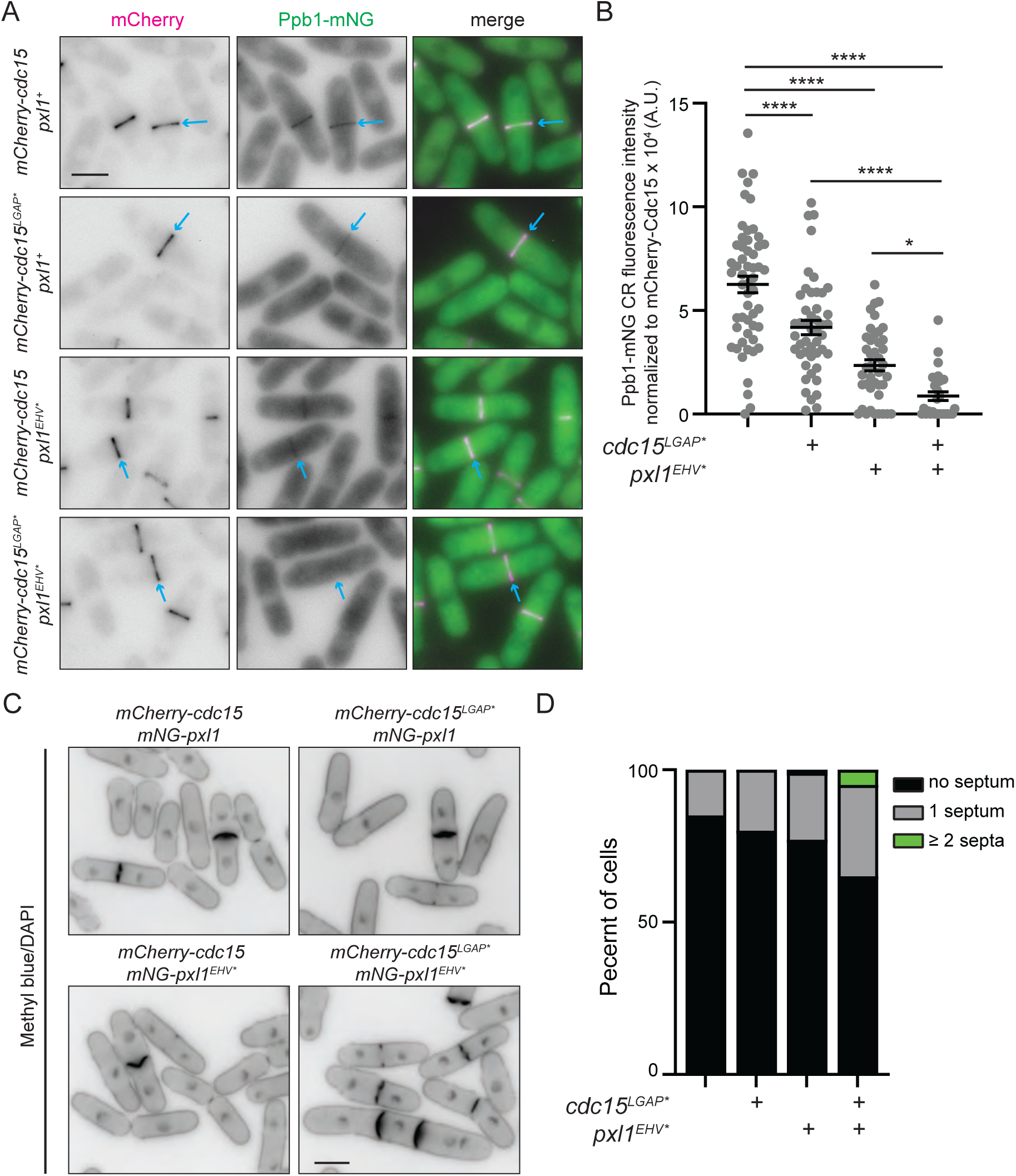
Cdc15 and Pxl1 cooperatively recruit calcineurin to the CR. **(A)** Live-cell images of cells expressing Ppb1-mNG and either mCherry-Cdc15 or mCherry-Cdc15^LGAP*^, in either wildtype *pxl1^+^* or *pxl1^EHV*^* cells. Blue arrows indicate examples of fully formed CRs that have not begun constriction. **(B)** Quantification of Ppb1-mNG fluorescence at fully formed CRs from A. *n* ≥ 41 from three biological replicates. \*\*\*\**p* > 0.0001, \**p* > 0.05; unpaired, two-tailed Student’s *t*-test. **(C)** Fixed-cell images of the indicated strains grown at 32°C in YE. Cells were stained with Methyl Blue and DAPI. **(D)** Quantification of septation phenotypes from C. *n* ≥ 317 cells per strain. Scale bars = 5 μm.

In accord with CN levels, we found that the mCherry-*cdc15^LGAP*^* and *pxl1^EHV*^* single mutants had largely normal septation profiles, with only *pxl1^EHV*^* showing a minor increase (∼1%) in cells with two or more septa (Fig 4C and 4D). However, the double mutant displayed cell separation defects similar to a complete loss of *ppb1* function with ∼6% of cells containing multiple septa (Fig 4C and 4D). These data indicate that both the Cdc15 LxVP-like motif and the Pxl1 LIM1 domain contribute independently and additively to CN recruitment and function at the CR.

### Identification of New CN Substrates During Cytokinesis

Cdc15 is a validated CN substrate at the CR (Martín-García et al., 2018; Snider et al., 2020). However, we reasoned that loss of Cdc15 dephosphorylation alone is unlikely to account for the cytokinesis defects observed in *ppb1Δ* cells. First, *cdc15* alleles harboring phospho-mimetic mutations do not phenocopy the cell separation defects seen in *ppb1Δ* cells (Roberts-Galbraith et al., 2010; Bhattacharjee et al., 2020, 2023). Also, we showed above that disrupting CN docking to Cdc15 (via the *mCherry-cdc15^LGAP*^* allele) does not impair cell separation (Fig 4C and 4D). We also combined *ppb1Δ* with *cdc15* alleles in which 11, 22, or 31 phospho-sites were mutated to alanine (Roberts-Galbraith et al., 2010; Bhattacharjee et al., 2020, 2023). However, all three double mutants (*ppb1Δ cdc15-11A*, *ppb1Δ cdc15-22A*, and *ppb1Δ cdc15-31A)* retained or exacerbated the cell separation defects seen in *ppb1Δ* alone (S2A and S2B Fig), the opposite of what would be expected if Cdc15 dephosphorylation was the only CN function during cytokinesis. These data strongly supported the idea that additional CN substrates, beyond Cdc15, are essential for successful cell division.

To identify such substrates, we took two approaches. First, because CN can bind substrates with a low affinity (Li et al., 2007, 2011; Roy et al., 2007), we used a proximity-based strategy to identify associated proteins in cells. For this, Ppb1 was tagged at its endogenous locus with the TurboID biotin ligase (BirA), and then biotinylated proteins were purified and identified by mass spectrometry (Larochelle et al., 2019). Because there are many native biotinylated *S. pombe* proteins (Tong, 2013; Larochelle et al., 2019), we conducted control experiments in which TurboID alone was expressed in cells or an unrelated protein (Ubx3) was tagged with TurboID and expressed in cells. We then considered proteins unique to, or greatly enriched in, the Ppb1-TurboID purification (S1 Table). Cnb1, the obligate Ppb1 binding partner, was a top hit and Cdc15 was also identified, validating the experimental approach.

Second, we performed a comparative phosphoproteomic analysis of synchronized, cytokinesis-arrested cells. *cps1-191* mutant cells arrest with fully assembled CN-containing CRs following a 3-hour temperature shift to 36°C (Liu et al., 1999). Thus, *cps1-191* arrested cells were treated with either 10 μg/mL of FK506 or vehicle control (DMSO) for 10 minutes, and the two phosphoproteomes were compared across three biological replicates using multiplexed quantitative mass spectrometry (Fig 5A). Reproducibility across replicates was high and, we identified 11,487 phosphopeptides from 2,523 proteins (Fig 5A). 179 phosphopeptides from 102 proteins were significantly upregulated (≥1.5-fold) in the FK506-treated samples (Fig 5A and S2 Table). Among the top phosphopeptides identified in our screen were seven from Cdc15 and three from Prz1, the two previously characterized CN substrates in *S. pombe* (Hirayama et al., 2003; Martín-García et al., 2018; Snider et al., 2020) (Fig 5A and S2 Table), validating the success of the approach.

**Fig 5.**
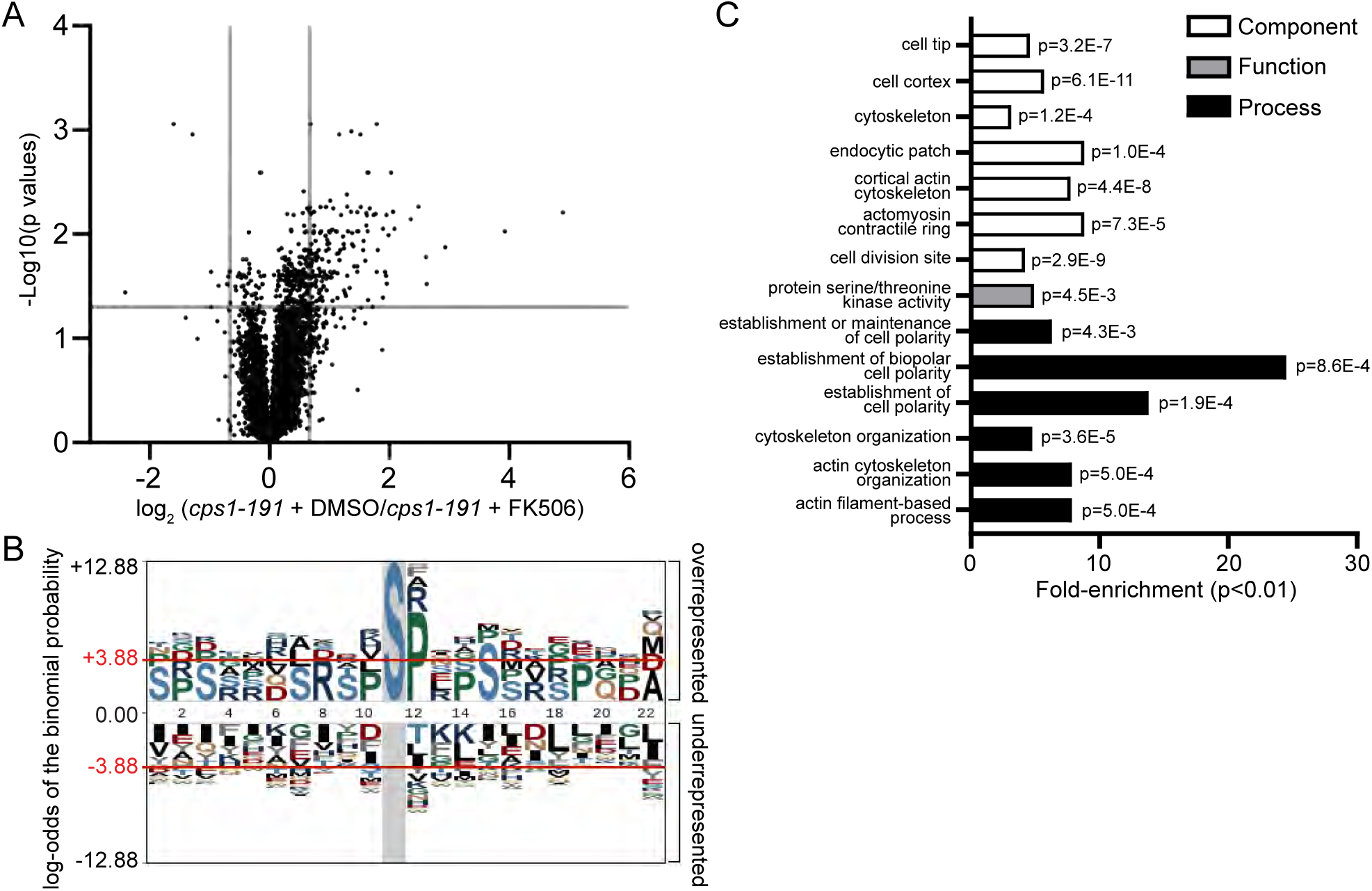
Phosphoproteomic analysis identifies candidate calcineurin substrates. (A) Volcano plot showing log₂(fold change) vs. -log₁₀(*p* value) for all quantified phosphosites in FK506- vs. DMSO-treated cells. Gray lines indicate significance thresholds: Benjamini-Hochberg adjusted *p* < 0.05 and fold change ≥ ±1.5. (B) Linear motif analysis of phosphopeptides upregulated ≥1.5-fold upon FK506 treatment. (C) GO term enrichment among proteins with FK506-sensitive phosphosites, relative to the *S. pombe* proteome. Bars indicate -log₁₀(*p* value).

Motif analysis of the FK506-enriched phosphopeptides revealed a strong preference for serines as the phosphorylated residue (O’Shea et al., 2013) (Fig 5B). Importantly, there was a significant enrichment of basic residues at the –3 position, a sequence feature that enhances CN-mediated dephosphorylation (Donella-Deana et al., 1994) (Fig 5B). In addition, many phosphosites contained a proline at the +1 position, suggestive of Cdk1 phosphorylation (Fig 5B). This is notable given that several known CN substrates are also Cdk1 targets (Arsenault et al., 2015; Ly and Cyert, 2017; Xiao et al., 2017) and Cdk1 is known to phosphorylate numerous CR components during mitosis (Wolfe et al., 2006; Roberts-Galbraith et al., 2010; Willet et al., 2018, 2021; Mangione et al., 2021; Akizuki et al., 2026).

Gene ontology analysis of the proteins harboring putative CN-regulated phosphosites revealed a strong enrichment for cellular components associated with cytokinesis (Paul D Thomas et al., 2022; Carme et al., 2026). These included proteins localized to the division site, actomyosin ring, cortical actin cytoskeleton, and related actin structures (Fig 5C). Functional enrichments were also observed for serine/threonine kinases, and the top biological process terms included regulation of the actin cytoskeleton and cell polarity, two processes tightly linked to cytokinesis (Fig 5C).

### Substrate validation

Besides Cdc15 and Prz1, 100 additional proteins contained at least one FK506-sensitive phosphosite, and of these, 58 were also identified in the TurboID experiment (S1 and S2 Tables). 18 of these proteins, like Cdc15, are known to localize at the cell division site (Carme et al., 2026). Of these, Nak1, Gef1, Rga7, Kin1, Sid2, and Cyk3 have known roles in cell division, either as scaffolding proteins or regulators (Sparks et al., 1999; Drewes and Nurse, 2003; Hirota et al., 2003; Huang et al., 2003; Pollard et al., 2012; Martín-García et al., 2014) (S3 Table). Thus, we first examined whether we could detect CN-dependent changes in their phosphorylation. Each was tagged endogenously with either the HA_3_, FLAG_3_ or Myc_13_ epitope in the *nda3-km311* mutant, which arrests in prometaphase prior to CN arriving at the CR (Hiraoka et al., 1984; Martín-García et al., 2018). Immunoprecipitates were left untreated or incubated with recombinant CN or λ phosphatase to determine if dephosphorylation occurred. Consistent with the compiled phosphosite data from proteome-wide screens (Carpy et al., 2014; Kettenbach et al., 2015; Swaffer et al., 2016, 2018; Lee et al., 2018; Tay et al., 2019; Halova et al., 2021; Mak et al., 2021), we found that all candidates were phosphorylated in the mitotic arrest (Fig 6A). We also determined that CN dephosphorylated all of these proteins (Fig 6A), validating our phosphoproteomics screen. While λ phosphatase eliminated all protein phosphorylation in this assay, CN partially dephosphorylated the candidate substrates (Fig 6A). We would not expect CN to target all phosphorylation sites on every substrate because other phosphatases likely contribute to the dephosphorylation of these CN substrates during mitotic exit.

**Fig. 6.**
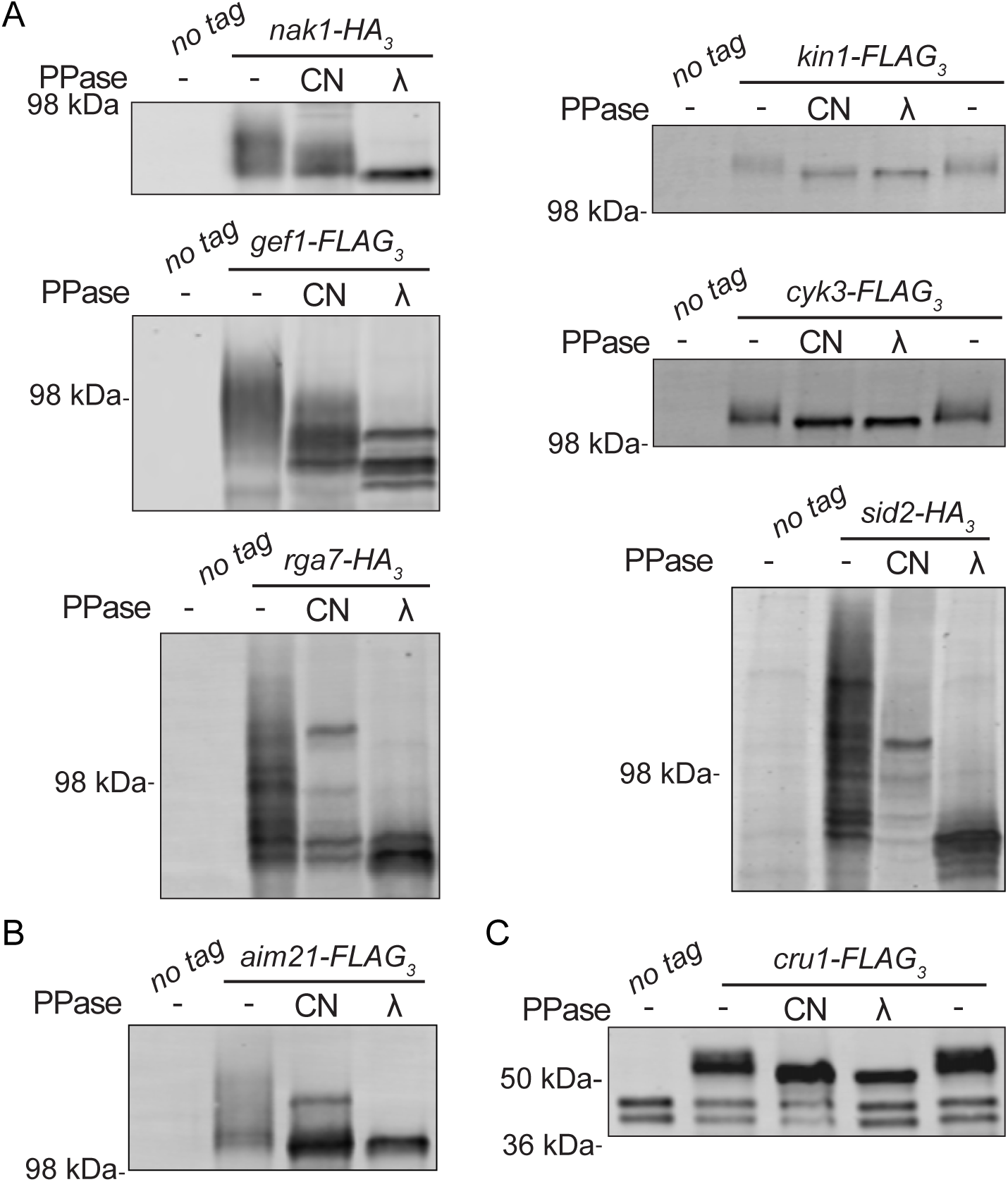
Validation of newly identified calcineurin substrates at the cell division site. (A-C) *nda3-km311* strains containing the indicated tagged proteins were arrested at 19°C for 6 h. Anti-FLAG, anti-HA or anti-Myc immunoprecipitates from the cell lysates were then treated with either vehicle control, recombinant CN, or lambda phosphatase (λ). Immunoprecipitates were then run on 6% SDS-PAGE Tris-Glycine gels (for Nak1, Kin1, Gef1, and Cyk3) or 6% SDS-PAGE Tris-Glycine gels containing 30-40 µM Phos-tag (for Rga7, Sid2, and Aim21). 10% NuPAGE Bis-Tris gels (NP0302) with MOPS SDS running buffer (NP0001, ThermoFisher Scientific) were used for Cru1.

In addition to the proteins described above, a phosphopeptide from Aim21 was a top hit in our phosphoprotomics screen (S2 Table). Aim21’s cellular localization has not been reported in *S. pombe* but based on its *S. cerevisiae* ortholog, Aim21 is expected to be a barbed end F-actin assembly factor (Shin et al., 2018) and thus possibly localized to the CR. We tagged Aim21 with mNG and found that, indeed, it localized to the CR as well as cortical actin patches in both asynchronously growing and *cps1-191*-arrested cells (S3A Fig). Aim21 is predicted to be disordered and proline-rich (9.6%) (Carme et al., 2026) and therefore it was not surprising that Aim21-FLAG_3_ migrated more slowly on SDS-PAGE gels than expected from its molecular mass (Fig 6B) (Theillet et al., 2013; Schramm et al., 2019). Like the other hits in our phosphoproteomics screen, Aim21-FLAG_3_ was hyperphosphorylated in the prometaphase arrest and partially dephosphorylated by CN (Fig 6B).

Phosphopeptides from an uncharacterized and unnamed protein, SPBC25B2.10, were also identified in the phosphoproteomics screen and an N-terminal fusion protein expressed from a plasmid was reported to localize to the cell division site (Matsuyama et al 2006). This protein is predicted to belong to the universal stress protein (USP) family present in bacteria and eukaryotes (Paul D. Thomas et al., 2022). We named this protein Cru1 for <u>C</u>alcineurin <u>R</u>egulated <u>U</u>niversal stress response protein <u>1</u>. We confirmed Cru1 localization to the division site with an endogenous C-terminal mNG tag (S3B Fig), and found that Cru1-FLAG_3_ was phosphorylated in a prometaphase arrest and partially dephosphorylated by CN (Fig 6C). Taken together, these results confirmed that there are multiple CN substrates at the cell division site in addition to Cdc15.

### Rcn1 is a CN substrate and negatively regulates CN CR localization

Phosphopeptides from a known CN inhibitor and substrate, Rcn1 (Kingsbury and Cunningham, 2000; Takasaki et al., 2024), were also enriched in our phosphoproteomics dataset (S2 Table). *S. pombe* Rcn1 is an ortholog of human RCAN1 (Regulator of Calcineurin 1), also referred to as calcipressin or Down syndrome critical region gene 1 (DSCR1) (Fuentes et al., 2000; Kingsbury and Cunningham, 2000; Rothermel et al., 2000; Takasaki et al., 2024; Carme et al., 2026). RCAN family proteins are well-established CN regulators; they can bind the phosphatase directly and modulate its catalytic activity (Görlach et al., 2000; Vega et al., 2002; Li et al., 2020; Ren et al., 2024b).

Consistent with Rcn1 being a direct and specific CN substrate, Rcn1-FLAG_3_ displayed FK506-dependent SDS-PAGE hyper-mobility (Fig 7A) and the higher mobility Rcn1-FLAG_3_ bands were collapsed upon treatment with CN or λ phosphatase (Fig 7B). As expected for a CN-specific substrate, Rcn1 phosphorylation state did not vary, as determined by Rcn1-FLAG_3_ SDS-PAGE mobility, in cells treated with other phosphatase inhibitors or in several other phosphatase deletion strains (S4A and S4B Fig) (Trautmann et al., 2001; Grallert et al., 2015). The two Rcn1 phosphosites identified in our screen, S97 and S101 (S2 Table), reside within a conserved CN pseudo-substrate motif, and are analogous to human RCAN1 residues S108 and S112, phosphorylation of which is implicated in controlling RCAN1-mediated catalytic inhibition of CN (Hilioti et al., 2004; Li et al., 2020). To validate that the sites we identified are the CN-targeted sites in *S. pombe* Rcn1, we mutated S97 and S101 to alanines (*rcn1-S97A,S101A*). Rcn1-S97A,S101A-FLAG_3_ was hypophorylated compared to wildtype Rcn1-FLAG_3_ *in vivo*. Indeed, the slower-migrating Rcn1-FLAG₃ species were largely absent from Rcn1-S97A,S101A-FLAG₃, and the gel mobility of Rcn1-S97A,S101A-FLAG₃ was the same as from CN-treated Rcn1-FLAG₃ (Fig 7B).

**Fig. 7.**
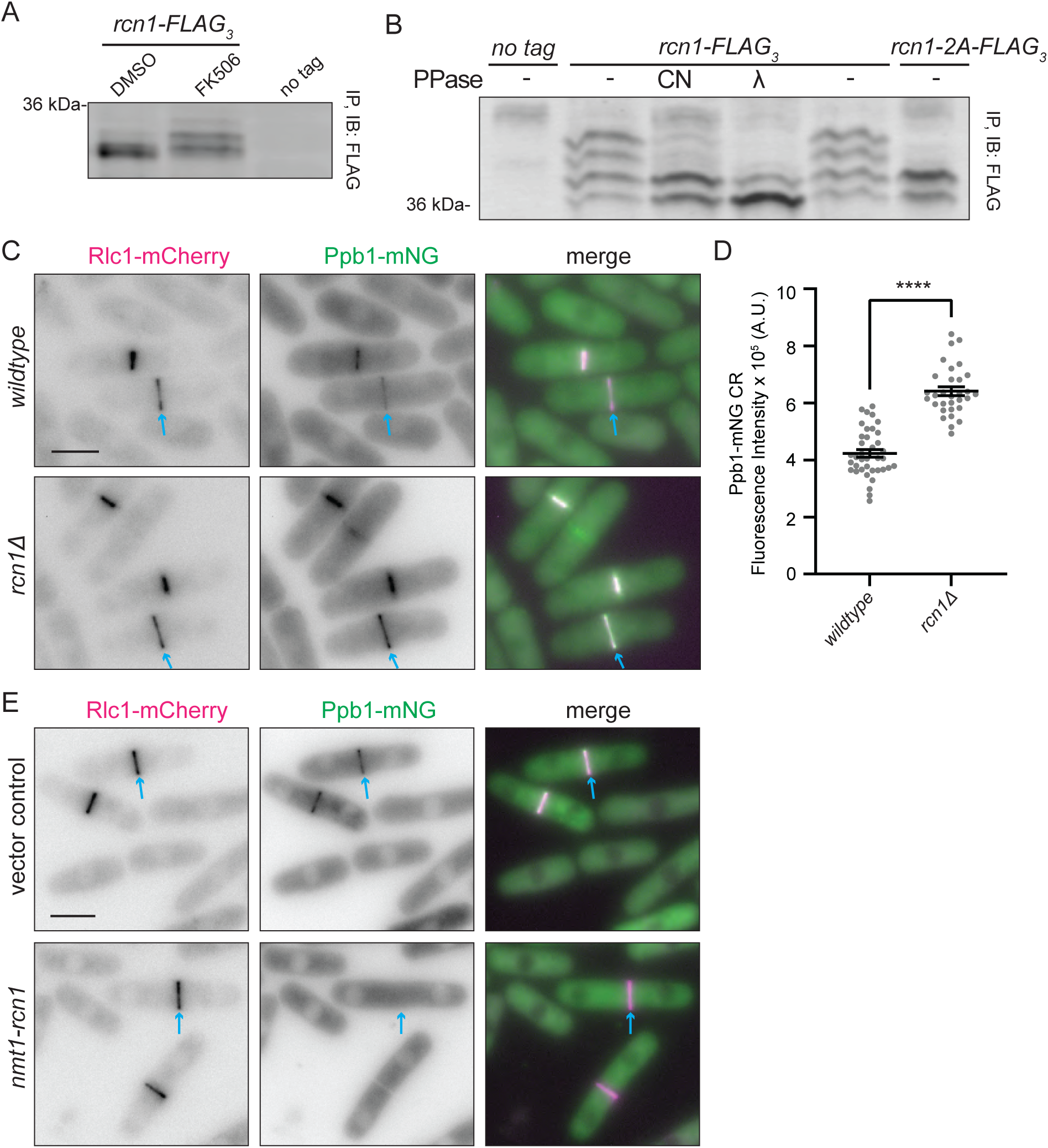
Rcn1 is a calcineurin substrate and negatively regulates calcineurin CR localization. **(A)** Rcn1-FLAG_3_ was immunoprecipitated from cells that had been treated with DMSO or 10 μg/ml FK506 for 30 min. The SDS-PAGE mobilities of the immunoprecipitates were then analyzed by immunoblotting. Phos-tag was incorporated into the gels to allow phosphorylation-induced mobility shifts to be more easily detected (see Materials and Methods). **(B)** Anti-FLAG immunoprecipitates from the indicated cell lysates were treated with either vehicle control, recombinant CN, lambda phosphatase (λ) or untreated. Immunoprecipitates were then run on 10% gels containing 30 µM Phos-tag. **(C)** Live-cell images of wildtype and *rcn1Δ* cells expressing Ppb1-mNG and Rlc1-mCherry. Blue arrows indicate examples of fully formed CRs that have not begun constriction. **(D)** Quantification of Ppb1-mNG intensity at fully formed CRs from A, normalized to whole-cell fluorescence. *n* ≥ 31 cells from two biological replicates. \*\*\*\**p* > 0.0001; unpaired, two-tailed Student’s *t*-test. **(E)** Live-cell images of cells overexpressing *rcn1^+^* from the *nmt1* promoter for 24 h. Ppb1-mNG and Rlc1-mCherry signals were assessed in cells carrying either the *rcn1^+^* plasmid or an empty vector (EV) control. Blue arrows indicate examples of fully formed CRs that have not begun constriction. Scale bars = 5 μm.

Despite detailed biochemical and structural characterization of the human RCAN1-CN complex (Vega et al., 2002; Li et al., 2020; Ren et al., 2024a), if and how Rcn1 regulates CN during cell division is not known. To examine this, we first determined its localization. Like other fungal RCAN homologs (Soriani et al., 2010; Harren et al., 2012), we found Rcn1-mNG to be diffusely cytoplasmic at all cell cycle stages (S4C Fig). This cytoplasmic localization was notable to us because a substantial portion of Ppb1-mNG signal is cytoplasmic, even during cytokinesis (Fig 1B). To investigate if Rcn1 regulates CN localization to the division site, we compared Ppb1-mNG CR localization in wildtype and *rcn1Δ* cells. Deletion of *rcn1Δ* led to a ∼50% increase in Ppb1-mNG at the CR (Fig 7C and D). Conversely, overexpression of *rcn1* abolished detectable Ppb1-mNG signal at the CR (Fig 7E), consistent with overexpression of *rcn1* mimicking the defects of CN loss-of-function mutants (Takasaki et al., 2024). We also found that Ppb1-mNG had increased CR abundance in *rcn1-S97A,S101A* cells compared to wildtype (Fig 8A and B). Thus, phosphorylation at S97 and S101 contributes to Rcn1-mediated restriction of CN accumulation at the CR. These data support a model in which Rcn1 sequesters CN in the cytoplasm, thereby limiting CN activity at the division site.

**Fig 8.**
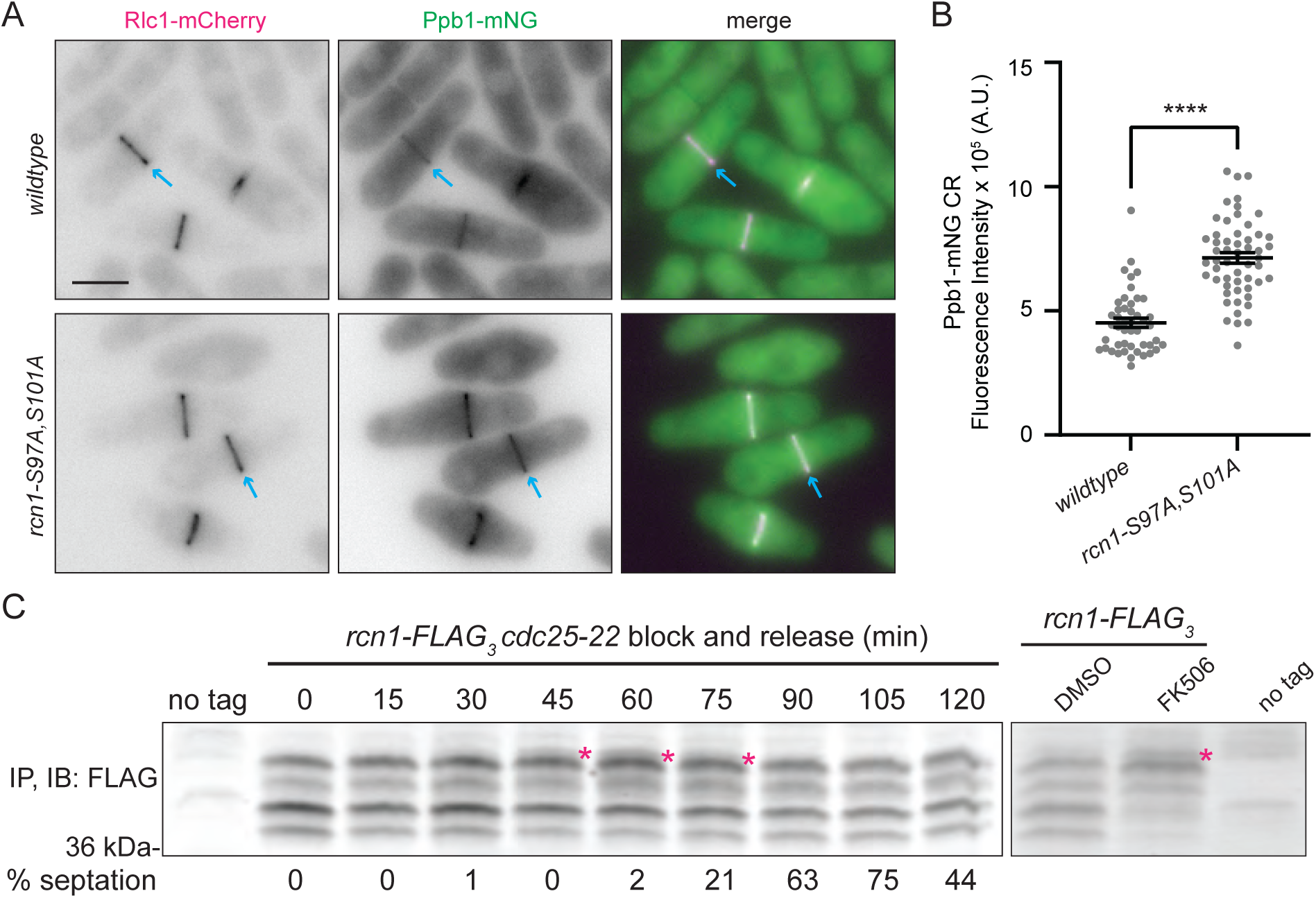
Rcn1 phosphorylation is cell-cycle regulated. **(A)** Live-cell imaging of wildtype and *rcn1-S97A,S101A* cells expressing Ppb1-mNG and Rlc1-mCherry. Blue arrows indicate example fully formed CRs that have not begun constriction. Scale bar = 5 μm. **(B)** Quantification of Ppb1-mNG intensity at fully formed CRs from G, normalized to whole-cell fluorescence. *n* ≥ 46 cells from three biological replicates. \*\*\*\**p* > 0.0001; unpaired, two-tailed Student’s *t*-test. **(C)** Left, cells were grown to mid-log phase, shifted to 36°C for 3 h, and then released to permissive temperature (25°C). Samples were collected at the indicated times and immunoprecipitated with Fab-TRAP beads. Samples were separated by 10% SDS-PAGE with 30 uM PhosTag and analyzed by immunoblot (IB) with an anti-FLAG antibody. Right, Rcn1-FLAG_3_ was immunoprecipitated from cells that had been treated with DMSO or 10 μg/ml FK506 for 30 min. The SDS-PAGE mobilities of the immunoprecipitates were then analyzed by immunoblotting. Phos-tag was incorporated into the gels to allow phosphorylation-induced mobility shifts to be more easily detected. This experiment is a biological replicate of Figure 7A.

To determine whether Rcn1 phosphorylation is cell cycle regulated, we monitored its phosphorylation state during synchronous mitotic progression. A hyperphosphorylated band of Rcn1-FLAG₃ became apparent during mitosis, peaking well before the peak of septation (Fig 8C). Interestingly, a similarly migrating hyperphosphorylated band was observed in FK506-treated cells (Fig 7A and 8C). Taken together, these results suggest that Rcn1 phosphorylation prior to septation contributes to limiting CN activity at the division site before cytokinesis begins.

## Discussion

In multiple organisms, CN localizes to the site of cell division (Juvvadi et al., 2008; Chircop et al., 2010; Kozubowski et al., 2011a; McDonald et al., 2017; Yadav et al., 2025). Here, we analyzed in detail how *S. pombe* CN is recruited to the division site, identified CN cell division substrates, and provided insight into how CN cell division site localization is regulated. Our findings support a multivalent docking mechanism in which Cdc15 and Pxl1 cooperatively recruit and position CN at the CR where it dephosphorylates multiple substrates to ensure non-defective cytokinesis.

Previous work established that both Pxl1 and Cdc15 are involved in recruiting CN to the division site and that Pxl1 and Cdc15 interact with each other (Martín-García et al., 2018; Mangione et al., 2019; Snider et al., 2020, 2022). Pxl1 binds Cdc15 through two interaction interfaces: an N-terminal motif that engages the cytoplasmic face of the Cdc15 F-BAR dimer (Fig 9, interface 1), and a central PXXP motif that interacts with the Cdc15 SH3 domain (Fig 9, interface 2) (Snider et al., 2020, 2022). We identified two additional interfaces on these proteins involved in CN division site recruitment: an LxVP-like motif within Cdc15 IDR2 that docks into the CN LxVP-binding pocket (Fig 9, interface 3), and a loop within the Pxl1 LIM1 domain that engages the CN PxIxIT-binding surface (Fig 9, interface 4). Together these interactions additively, rather than redundantly, contribute to CN CR recruitment. This interaction model is consistent with the CR architecture defined by super-resolution microscopy, which places the Cdc15 F-BAR domain within the membrane-proximal layer, CN within the intermediate layer, and the Cdc15 SH3 domains within the most membrane-distal CR layer (Laplante et al., 2016; McDonald et al., 2017). The positioning of the Cdc15 N- and C-termini in distinct CR layers combined with CN-dependent dephosphorylation of the Cdc15 IDR are together proposed to help maintain Cdc15 in an open conformation to promote Cdc15 protein partner binding, oligomerization, and membrane binding with high avidity (Roberts-Galbraith et al., 2010; McDonald et al., 2015, 2017; Bhattacharjee et al., 2020, 2023; Snider et al., 2020). CN positioned in the CR’s intermediate layer is physically near not only to Cdc15, but also additional substrates whose dephosphorylation we propose is critical for the proper execution of cytokinesis (Martín-García et al., 2018; Snider et al., 2020, 2022).

**Fig 9.**
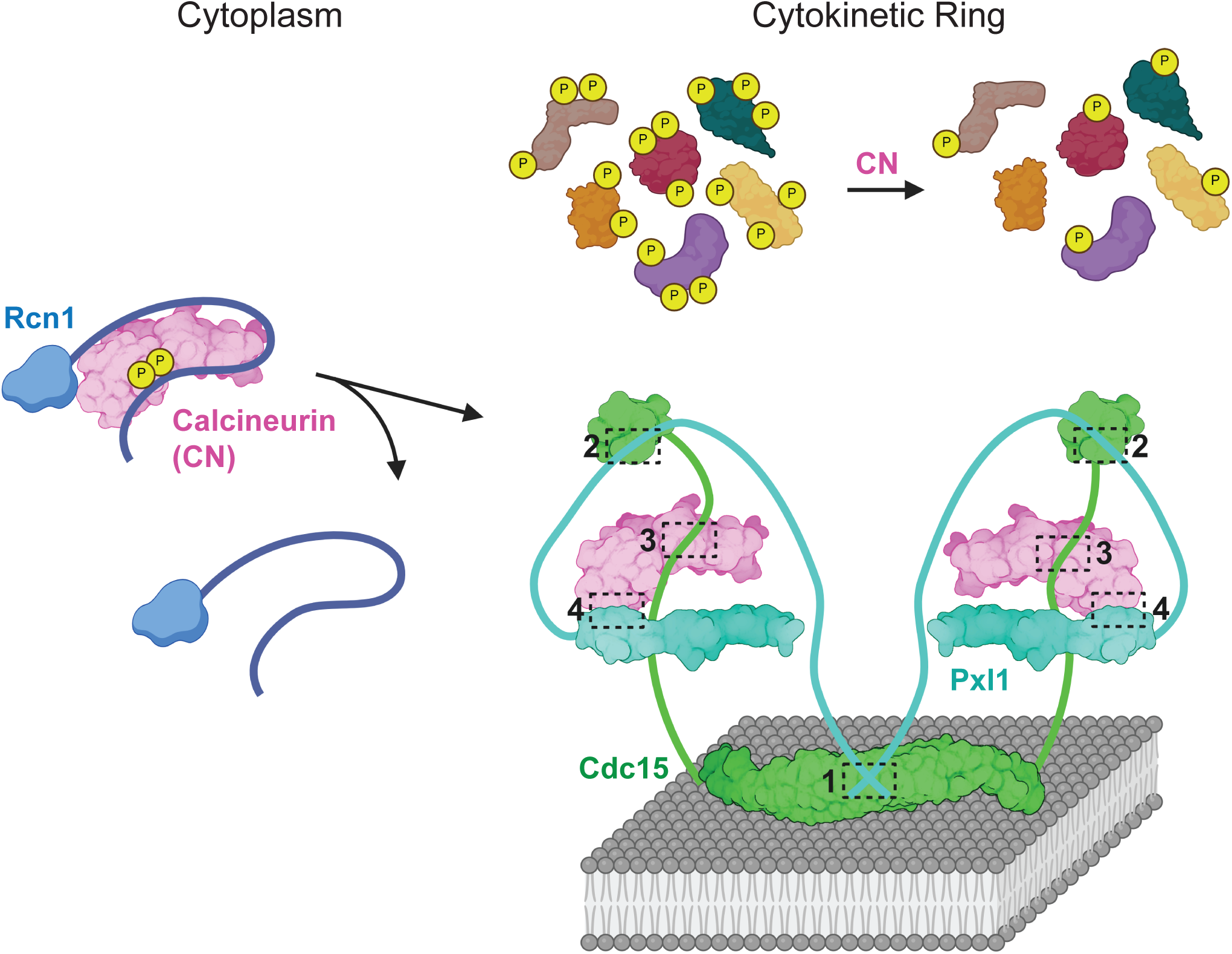
Model for calcineurin localization to the cytokinetic ring and dephosphorylation of cytokinetic substrates. When CN (pink) is bound to phosphorylated (yellow) Rcn1 (blue) in the cytoplasm, CN is kept in an inhibited state. Upon dephosphorylation of Rcn1, the inhibitory complex is relieved, and CN is permitted to localize to the CR. At the CR, four interaction interfaces link Cdc15 (green), CN (pink), and Pxl1 (cyan). Interface 1: The cytoplasmic face of the Cdc15 F-BAR domain, bound to the plasma membrane, binds a motif at the extreme N-terminus of Pxl1 (Snider et al., 2020). Interface 2: The Cdc15 SH3 domain interacts with a PXXP motif in the Pxl1 N-terminus (Snider et al., 2020). Interface 3: A conserved LxVP-like motif in the Cdc15 IDR2 region binds the LxVP docking site of Ppb1. Interface 4: A loop within the Pxl1 LIM1 domain engages the PxIxIT docking site of Ppb1. CN partially dephosphorylates a cohort of substrates at the CR to promote proper completion of cytokinesis.

This multivalent, SLiM-mediated tethering strategy parallels principles established for CN localization in other contexts. For example, in mammalian cells, the scaffold protein AKAP79 recruits CN to the plasma membrane through a PxIxIT-like motif (Li et al., 2012). However, the functional consequences of AKAP79-mediated CN recruitment remain debated. Some studies suggest that CN bound to AKAP79 is maintained in an inhibited state (Taigen et al., 2000; Dell’Acqua et al., 2002), whereas others propose that AKAP79 concentrates CN near its substrate, NFAT, thereby promoting efficient dephosphorylation and activation of NFAT (Li et al., 2012). We propose that the primary function of Cdc15- and Pxl1-mediated docking is to concentrate CN near its cytokinetic substrates. Disruption of Cdc15- and Pxl1-mediated CN docking nearly eliminated CN at the CR and produced cytokinetic defects that resembled CN loss-of-function phenotypes, indicating that localization of CN to the division site is required for efficient substrate dephosphorylation during cytokinesis. We note that it is intriguing that the LxVP- and PxIxIT-binding surfaces appear to mediate interactions with both CN scaffolds and substrates in multiple contexts (Taigen et al., 2000; Dell’Acqua et al., 2002; Li et al., 2011, 2012). How these interactions are coordinated to promote efficient substrate dephosphorylation while maintaining stable recruitment of CN to sites of action remains an important unanswered question.

Given the significant cell division defects of cells lacking CN function (Yoshida et al., 1994; Chircop et al., 2010; Yadav et al., 2025), a central motivation for this study was to identify CN substrates involved specifically in this process. Combining TurboID proximity labeling with cell cycle stage-specific quantitative phosphoproteomics revealed a cohort of candidate CN substrates enriched at the cell division site. We experimentally confirmed CN-dependent dephosphorylation of six proteins with established roles in cell division (Nak1, Gef1, Rga7, Kin1, Sid2, and Cyk3) (Sparks et al., 1999; Hirota et al., 2003; Huang et al., 2003; Pollard et al., 2012; Cadou et al., 2013; Martín-García et al., 2014). Interestingly, Cyk3 is a component of the ingression progression complex in budding yeasts and it is notable that another component of this complex was identified as a CN substrate in *Cryptococcus deneoformans* (Yadav et al., 2025). Our results also implicate Aim21, a putative barbed-end actin assembly factor that we showed localizes to the CR, and Cru1, a previously uncharacterized universal stress protein family member that we confirmed also localizes to the cell division site, as having roles in cell division, an implication that can inform future research efforts. Although our work focused on validating CR-localized substrates, our dataset includes many other proteins involved in processes known to influence cytokinesis such as those involved in intracellular trafficking as well as membrane and ion homeostasis (Carme et al., 2026) (S2 Table). Together with Cdc15, these findings expand the known cytokinetic CN substrate network and provide a framework for understanding how CN regulates cell division through the concerted dephosphorylation of multiple cellular targets.

The majority of CN substrates we validated were highly phosphorylated during mitosis and only partially dephosphorylated by recombinant CN, suggesting that CN targets only a subset of phosphorylation sites on each protein. This suggests that CN functions in concert with other mitotic phosphatases to coordinate substrate dephosphorylation during mitotic exit. Consistent with this idea, *ppb1Δ* exhibits synthetic sick or lethal genetic interactions with mutations affecting other phosphatases (Yoshida et al., 1994; Tanabe et al., 2001). An additional layer of complexity is that at least three CN substrates (Nak1, Kin1 and Sid2) are themselves protein kinases, suggesting that CN-mediated dephosphorylation may reshape phosphorylation networks beyond its direct substrates.

Our findings suggest that in *S. pombe*, Rcn1 negatively regulates CN localization to the cell division site, as deletion of *rcn1* increased CN abundance at the CR, whereas overproduction diminished CN accumulation at the CR. Consistent with previous models in which Rcn1/RCAN proteins participate in negative feedback regulation of CN (Kingsbury and Cunningham, 2000; Hilioti et al., 2004; Mehta et al., 2009), our data support a model in which phosphorylated Rcn1 restricts CN activity not only through catalytic inhibition but also by limiting the pool of CN available for recruitment to the division site. In this model, Rcn1 could sequester CN in the cytoplasm, thereby preventing excessive accumulation of active CN at the CR. We further observed that Rcn1 becomes hyperphosphorylated prior to septation, raising the possibility that this regulatory mechanism is modulated with cell-cycle progression. Such regulation could provide a temporal layer of control over CN availability, helping to restrict when and how much CN is recruited to the division site.

Collectively, our findings support a model in which CN is regulated during cell division through multivalent scaffold interactions that position the enzyme, and feedback from an RCAN-family inhibitor that controls its availability. These combined inputs allow for coordinated dephosphorylation of a diverse set of cytokinetic substrates, ensuring proper completion of cytokinesis.

## Materials and Methods

### Yeast methods

*Schizosaccharomyces pombe* strains used in this study are listed in Table S4. Cells were cultured in yeast extract (YE) medium under standard conditions unless otherwise noted (Forsburg and Rhind, 2006). Gene tagging at the 3′ end of open reading frames (ORFs) was performed using PCR-based homologous recombination with pFA6a-based cassettes encoding FLAG₃:*hphMX6*, HA_3_:*kanMX6*, mCherry:*kanMX6*, mNG:*kanMX6*, mNG:*hphMX6* or V5-TurboID*:kanMX6* turbo ID (Wach et al., 1994; Bähler et al., 1998). Correct integration of tags or truncations was confirmed by colony PCR and/or fluorescence microscopy. All fusion proteins were expressed from their endogenous promoters at their native chromosomal loci unless otherwise indicated.

Transformations were carried out using the lithium acetate method (Keeney and Boeke, 1994). Gene-tagged strains were crossed into relevant genetic backgrounds using standard *S. pombe* mating, sporulation, and tetrad dissection protocols (Forsburg and Rhind, 2006).

Various *ppb1* alleles were constructed by cloning the *ppb1⁺* ORF along with 500 bp of upstream and downstream untranslated regions into the pIRT2 vector at the *PstI* and *BamHI* sites using Gibson Assembly. The *rcn1* mutant was constructed by cloning the *rcn1* ORF with 300 bp of upstream and downstream untranslated regions into the pIRT2 vector at the *BamHI* site using Gibson Assembly. Site-directed mutagenesis was used to introduce specific point mutations into the genes, and these mutations were confirmed by DNA sequencing.

To integrate mutants at the endogenous loci, plasmids were transformed into *ppb1Δ::ura4⁺* or *rcn1::ura4^+^* cells. Transformants were selected on Edinburgh minimal medium (EMM) supplemented with adenine. These colonies were then grown in liquid YE medium for 24 hours, and the cells were plated on YE containing 1.5 mg/mL 5-fluoroorotic acid (5-FOA; United States Biological, F5050) to select for recombinants that had lost the *ura4*⁺ marker. Correct integration was verified by colony PCR and DNA sequencing.

### Microscopy

For live-cell imaging, *S. pombe* cultures were grown in YE medium and grown at 25°C. Images in Figure 1B were acquired using a DeltaVision imaging system (Leica Microsystems) built on an Olympus IX71 inverted microscope equipped with a 60× NA 1.42 Plan Apochromat and 100× NA 1.40 U-Plan S-Apochromat objective. The system includes standard and live-cell filter wheels, a pco.edge 4.2 sCMOS camera, and softWoRx software for image acquisition.

Images for Figures 1D and F, 2A and C, 4A and C, 7C, E and 8A, as well as Figures S1, S2A, S3, and S4C were acquired using a Zeiss Axio Observer inverted epifluorescence microscope with a 63× oil immersion objective (NA 1.46). Images were captured using a Zeiss Axiocam 503 monochrome camera and processed with ZEN 3.0 (Blue edition) software.

Live-cell time-lapse imaging for Figure 2E was performed using a Leica Thunder Imager system with a DMi8 inverted microscope and a 63× Plan Apochromat oil immersion objective (NA 1.40). A Leica K8 sCMOS camera, standard excitation/emission filters, and LED light source were used for fluorescence acquisition, controlled via Leica Application Suite X (LAS X) software. A CellASIC ONIX microfluidics system (Millipore Sigma) with Y04C microfluidic plates (Millipore Sigma, Y04C-02-5PK) was used for cell immobilization. Cells were loaded for 10 s at 8 psi, and YE medium was perfused continuously at 5 psi during imaging. Z-stacks were acquired at 0.5 µm intervals over 4.5 µm every 2 min.

All images used for quantification were non-deconvolved sum projections. Background correction was performed by measuring the average pixel intensity in a cell-free region, multiplying this value by the area of the region of interest (ROI), and subtracting the result from the raw ROI intensity. Image quantification and fluorescence intensity measurements were performed using Fiji/ImageJ (available at https://fiji.sc) (Schindelin et al., 2012).

### Recombinant protein expression and purification

CN was produced by co-expressing the catalytic subunit Ppb1 (tagged with His₆ or His₆-SPOT), the regulatory subunit Cnb1, and calmodulin (Cam1) in *Escherichia coli* Rosetta2(DE3)pLysS cells, as previously described (Snider et al., 2020). Cultures were grown in Terrific Broth (23.6 g/L Yeast Extract, 11.8 g/L tryptone, 9.4 g/L K_2_HPO_4_, 2.2 g/L KH_2_PO_4_, and 4 mL/L glycerol) to log phase, induced with 0.5 mM isopropyl β-D-1-thiogalactopyranoside (IPTG), and incubated overnight at 20°C. Cells were harvested by centrifugation (6,000 × g, 10 min, 4°C), lysed by sonication (3 × 30 s, 30 s pause, 15 W; Sonic Dismembrator Model F60, Fisher Scientific) in lysis buffer (50 mM Tris-HCl, pH 7.4, 150 mM NaCl, 0.1% NP-40), and supplemented with cOmplete EDTA-free protease inhibitor cocktail (Roche, 05056489001). Clarified lysate was incubated with Ni²⁺-NTA resin (Cytiva, 17371201), washed extensively with lysis buffer, and eluted with buffer containing 200 mM imidazole. Eluted protein was dialyzed into storage buffer (50 mM Tris-HCl, pH 7.4, 150 mM NaCl, 0.1% NP-40, 2 mM CaCl₂, 10 mM MgCl₂).

Cdc15(441-927) was expressed as a glutathione S-transferase (GST) fusion protein in Rosetta2(DE3)pLysS cells. Cultures were grown in Terrific Broth, induced with 0.5 mM IPTG at 20°C overnight, and harvested by centrifugation. Cells were lysed by sonication in phosphate-buffered lysis buffer (4.3 mM NaHPO₄, 137 mM NaCl, 2.7 mM KCl, 1 mM DTT, 1 mM PMSF, 1.3 mM benzamidine), and the cleared lysate was incubated with GST-bind resin (Millipore Sigma, 70541). Resin was washed thoroughly with lysis buffer. GST alone was purified in parallel and used as a negative control in binding assays.

MBP-V5-Pxl1 was expressed in the presence of 150 µM ZnCl₂ to support zinc finger stability. Cells were lysed in buffer (20 mM Tris-HCl, pH 7.4, 150 mM NaCl, 1 mM DTT) containing 200 µg/mL lysozyme (Sigma-Aldrich, L6876), cOmplete EDTA-free protease inhibitor cocktail, and 0.1% NP-40 (US Biologicals, N3500). Cell suspensions were incubated on ice for 20 min with gentle agitation, followed by sonication. Lysates were clarified by centrifugation (10,000-13,000 rpm, 15-30 min), and the supernatant was incubated with amylose resin (New England Biolabs, E8021L) for 2 h at 4°C with nutation. Resin was washed three times with lysis buffer and resuspended as a 50% slurry. For elution, V5-Pxl1 was cleaved from MBP by incubating resin-bound protein with 2 units of PreScission Protease (Cytiva, 27-0843-01) overnight at 4°C. Protein concentration was determined via SDS-PAGE and Coomassie Brilliant Blue G staining (Sigma-Aldrich, B0770), using bovine serum albumin (BSA; Sigma-Aldrich, A9418) as a standard.

### In vitro binding assays

All *in vitro* binding assays were performed in a modified NP-40 lysis buffer (50 mM Tris-HCl, pH 7.4, 150 mM NaCl, 0.1% NP-40) lacking EDTA and supplemented with 1 mM CaCl₂ to support CN activity. For Cdc15-CN binding assays, 4 µg of bead-bound GST or GST-Cdc15 fusion protein and 5 µg of purified CN complex (Ppb1, Cnb1, and Cam1) were incubated in 200 µL of modified NP-40 lysis buffer at 4°C for 30 min with gentle rotation. Following incubation, beads were washed four times with 500 µL of binding buffer to remove unbound proteins. Beads were resuspended in 40 µL of 2× SDS sample buffer, and 20 µL of each reaction was resolved on a 4-12% NuPAGE gel (Thermo Fisher Scientific, NP0321). Gels were stained with Coomassie Brilliant Blue and imaged using an Odyssey CLx imaging system (LI-COR Biosciences).

For Pxl1-CN binding assays, 5 µg of bead-bound CN complex (His₆-SPOT-Ppb1, Cnb1, and Cam1) was incubated with 8-10 µg of V5-tagged Pxl1 (either full-length or truncation variants) purified from *E. coli* lysates. Binding reactions were carried out in 200 µL of Ca²⁺-supplemented NP-40 buffer at 4°C for 1 h with gentle nutation. Beads were washed four times with 500 µL of binding buffer, resuspended in 40 µL of 2× SDS sample buffer, and 20 µL of each sample was resolved by SDS–PAGE. Proteins were transferred to PVDF membranes and probed with anti-V5 and anti-His primary antibodies. Blots were developed and imaged on an Odyssey CLx system.

### Immunoprecipitations and phosphatase assays

For CN and λ-phosphatase assays, pellets (30 OD) collected from asynchronous or mitotically arrested cells were flash frozen then lysed by glass bead disruption in NP-40 buffer (6 mM Na_2_HPO_4_, 4 mM NaH_2_PO_4_, 1% NP-40, 150 mM NaCl, 50 mM NaF, 4 μg/mL leupeptin, 0.1 mM Na_3_VO_4_)(Gould et al., 1991). Lysates were denatured by boiling at 95°C for 1.5 min in 300 µL of SDS lysis buffer (6 mM Na_2_HPO_4_, 4 mM NaH_2_PO_4_, 0.5% SDS, 1 mM EDTA, 50 mM NaF, 1 mM DTT, 4 μg/mL leupeptin, 0.1 mM Na3VO4), followed by extraction with 800 µL of NP-40 buffer with the addition of 1 mM PMSF, 1.3 mM benzamidine, 0.2 mM diisopropyl fluorophosphate (DIFP), and Roche phosphatase and protease inhibitor cocktail. Proteins were immunoprecipitated from denatured lysate using 4 µg α-HA (Vanderbilt Antibody and Protein Resource, Anti-HA 12CA5, RRID: AB_2923038) and 80 µL 50% slurry of protein G Sepharose (GE Healthcare, P3296), or 12.5 µL 50% slurry of Fab trap beads (Proteintech, ffa). For CN phosphatase assays, beads were washed twice with NP-40 buffer then washed three times with calcineurin buffer (50 mM Tris pH 7.4, 150 mM NaCl, 0.1% NP-40, 2 mM CaCl_2_, 10 mM MgCl_2,_ 1 mM DTT). One half was treated with 20 μg purified CN and the other half was treated with a same volume of CN buffer for control. For λ-phosphatase assays, the λ-phosphatase buffer (50 mM HEPES, 100 mM NaCl, 2 mM DTT, 0.01% Brij 35, pH 7.5) was used instead and 1 µL λ-phosphatase (NEB, P0753S) or λ-phosphatase storage buffer as control was added in the presence of 1 mM MnCl_2_.The reactions were incubated for 35 min at 30°C with shaking. Samples were then resolved by SDS-PAGE [30-50 µM Phostag (Wako Chemicals, 304-93521) were included for better detection of protein phosphorylation], transferred to PVDF. FLAG was detected with a monoclonal antibody M2 (Sigma, F-1804, RRID: AB_262044) at a 1:1,000 dilution. HA was detected with a monoclonal antibody 12CA5 (Vanderbilt Antibody and Protein Resource, Anti-HA 12CA5, RRID: AB_2923038) at a 1:1000 dilution, ɑ-tubulin was detected with a monoclonal antibody (Sigma-Aldrich, T6557, RRID: AB_477584) at a 1:10,000 dilution, His was detected with monoclonal antibody to penta-his (Invitrogen, P-21315, RRID: AB_2539819) and V5 was detected with a monoclonal antibody (GenScript, A01724, RRID: AB_2622216) at 1 μg/mL final concentration. Primary antibodies were detected with secondary antibodies coupled to Alexa Fluor 680 goat anti-mouse (LICOR Biosciences, 926-68020, RRID: AB_10706161) or goat anti-rabbit (LI-COR Biosciences, 926-68021, RRID: AB_10706309) or IRDye800 goat anti-mouse (LI-COR Biosciences, 926-32210, RRID: AB_621842) or goat anti-rabbit (LI-COR Biosciences, 926-32211, RRID: AB_621843) all at a 1:10,000 dilution and visualized using an Odyssey Infrared Imaging System (LI-COR Biosciences).

For *nda3-KM311* cells treated with FK506 (LC Laboratories, F-4900) or DMSO, the cells were grown at 32°C and then shifted to 19°C for 6 h. Next, the cells were transferred back to 32°C and DMSO or 10 ng/μL of FK506 was added for 30 min before 30 OD pellets were collected and snap frozen. Pellets were then processed the same as described in the above paragraph. Protein samples were resolved by SDS-PAGE in the presence of 25 or 50 μM Phos-tag acrylamide per the manufacturer’s protocol. For Rcn1-FLAG_3_, immunoprecipitations were treated with 100 nM calyculin A (Cell Signaling, 9902) or 1 μM okadaic acid (Sigma-Aldrich, 495604) prior to running samples with SDS-PAGE.

Cells were arrested in G2 with the *cdc25-22* allele by being shifted to the restrictive temperature (36°C) for 3 h and then released back into the cell cycle by shifting to the permissive temperature (25°C). 20 OD cell pellets were collected at the time of release (0 min) and at all subsequent time points. Additionally, an aliquot of cells was fixed in ice-cold 70% ethanol to determine the percent of septation at each time point (see Microscopy and image analysis section).

### Mass spectrometry

For the Turbo-ID experiments, mass spectrometric analysis was performed as described previously (Beckley et al., 2015; Willet et al., 2024). Streptavidin beads from TurboID purifications were washed with Tris-urea buffer (100 mM Tris-HCl, pH 8.5, 2 M urea) three times. Proteins were reduced with 3 mM Tris(2-carboxyethyl)phosphine hydrochloride, alkylated with 10 mM chloroacetamide, and digested with trypsin (1 μg of Trypsin Gold, Promega, V5280) at 37°C overnight with shaking. The digest supernatant and washes were combined, concentrated and desalted with a Pierce C18 spin column (Thermo Fisher Scientific, 89870). Peptides were resuspended in 0.1% formic acid and loaded onto a 26 cm column [consisting of 3 cm of 5 µm Jupiter C-18 (Phenomenex), 3 cm of 5 µm SCX (Phenomenex) and 20 cm of 3 µm Jupiter C-18 in 100 µm fused silica capillary tubing] with a pressure cell and then separated and analyzed by three-phase multidimensional protein identification technology (MudPIT) on a Velos linear trap quadrupole (LTQ) mass spectrometer (Thermo Fisher Scientific) coupled to a nanoHPLC (NanoAcquity; Waters Corporation). The NanoAcquity autosampler was used to inject 2 μl of varying concentrations of ammonium acetate (0, 10, 25, 50, 100, 200, 300, 400, 600, 800, 1000 and 5000 mM) for 11 salt elution steps. Each injection was followed by elution of peptides with a 2–40% acetonitrile gradient (60 min) except the first and last injections, in which a 2–90% acetonitrile gradient was used. One full precursor mass MS scan (400-2000 mass-to-charge ratio) and five tandem MS (MS2) scans of the most abundant ions detected in the precursor MS scan under dynamic exclusion were performed. Ions with a neutral loss of 98 Da (singly charged), 49 Da (doubly charged), or 32.7 Da (triply charged) from the parent ions during MS2 were subjected to MS3 fragmentation.

RAW files containing more than 20 peaks were converted to DTA files using Scansifter software (Ma et al., 2011) (v2.1.25). Each DTA file was searched using the SEQUEST algorithm (Thermo Fisher Scientific; version 27, rev. 12). SEQUEST was set up to search the pombe_contams_20151012_20151012_rev database (downloaded from PomBase in October 2015 with common contaminants added and all sequences reversed, 10390 entries in total). Variable modifications (C+57, M+16, [STY]+80, [STY]−18), strict tryptic cleavage, <10 missed cleavages, fragment mass tolerance: 0.00 Da (this results in 0.5 Da tolerance in SEQUEST), and parent mass tolerance: 2.5 Da were allowed. Peptide identifications were assembled and filtered in Scaffold (v4.7.5, Proteome Software) using the following criteria: minimum of 99% protein identification probability; minimum of two unique peptides; minimum of 95% peptide identification probability. These filtering criteria were used to achieve false discovery rates less than 1%.

Sample preparation for quantitative phosphoproteomics was performed following the SL-TMT protocol (Navarrete-Perea et al., 2018). Logarithmically growing *S. pombe cps1-191* cells were grown up at 25°C and shifted to 36°C for 3 h. DMSO or 10 ng/μL of FK506 was added to the cells and incubated at 36°C for 5 min before sample collection. 15 OD cell pellets were collected by centrifugation and snap frozen in liquid nitrogen. Cell pellets were lysed by bead-beating in 8 M urea complemented with protease and phosphatase inhibitors. After lysis, the protein extracts were centrifuged, and the supernatant was obtained. Samples were reduced using 5 mM TCEP for 30 min, alkylated with 10 mM iodoacetamide for 30 min, and the excess of iodoacetamide was quenched using DTT. After protein quantification, 200 μg of protein were chloroform-methanol precipitated and reconstituted in 200 μL of 200 mM EPPS pH 8.5. Protein was digested using LysC overnight at room temperature followed by trypsin for 6 h at 37°C, both at a 100:1 protein-to-protease ratio. After digestion, the samples were labeled using the TMTpro16 reagents (Thermo Fisher Scientific, A44520) for 60 min, the reactions were quenched using hydroxylamine (final concentration of 0.3% v/v) for 20 min. After label check, the samples were combined equally and desalted. Phosphopeptides were enriched using the Pierce High-Select Fe-NTA Phosphopeptide Enrichment kit (Thermo Fisher Scientific, A32992) following the manufacturer’s instructions. The phosphopeptides were eluted in a tube containing 100 μL of 10% formic acid and dried in a vacuum centrifuge. The unbound fraction was retained for whole proteome analysis. The phosphopeptides and whole proteome were fractionated using the Pierce High pH Reversed-Phase Peptide Fractionation Kit (Thermo Fisher Scientific, 84868) following the manufacturer’s instructions. The peptides were eluted using the following ACN concentrations: 7.5, 10, 12.5, 15, 17.5, 20, 22.5, 25, and 50% ACN, then the samples were combined intro 6 final fractions.

Mass spectrometry data were collected on an Orbitrap Eclipse mass spectrometer coupled to a Proxeon NanoLC-1200 UHPLC. The peptides were separated using a 100 μm capillary column packed with ≈30 cm of Accucore 150 resin (2.6 μm, 150 Å; Thermo Fisher Scientific). Samples were analyzed using FAIMS/hrMS2 following our optimized workflow for multiplexed proteomics and phosphoproteomics analysis (Schweppe et al., 2020b, 2020a). The global proteome fractions were analyzed with three CVs (CV = −40 V, −60 V and −80 V) and a 90-min gradient of 6% to 30% B, while the phosphorylated peptides were analyzed with three CVs (CV = −40 V, −60 V and −80 V) and a 150-min gradient of 6% to 30% B.

Raw files were first converted to mzXML. Database searching included all *Schizosaccharomyces pombe* entries from UniProt (downloaded August 2023). The database was concatenated with one composed of all protein sequences in the reversed order and a list of common contaminant proteins was also included. Searches were performed using a 50 ppm precursor ion tolerance and 0.03 Da (high-resolution MS2) product ion tolerance (Beausoleil et al., 2006; Elias and Gygi, 2007; Huttlin et al., 2010). TMTpro on lysine residues and peptide N termini (+304.2071 Da) and carbamidomethyleneation of cysteine residues (+57.0215 Da) were set as static modifications (except when testing for labeling efficiency, when the TMTpro modifications are set to variable). Oxidation of methionine residues (+15.9949 Da) was set as a variable modification. For phosphopeptide analysis, +79.9663 Da was set as a variable modification on serine, threonine, and tyrosine residues. The mass spectrometry proteomics data have been deposited to the ProteomeXchange Consortium via the PRIDE (Perez-Riverol et al., 2019) partner repository with the dataset identifier PXD083328.

*t* tests were used to compare each protein or phosphorylation site measurement, and a Benjamini-Hochberg multiple testing correction was applied. Statistical analysis and volcano plots were generated in Graph Pad Prism v8. Linear substrate motif analysis was performed on phosphopeptides with *P* < 0.01 using pLogo (O’Shea et al., 2013). For gene ontology analysis the phosphorylation sites from all phosphopeptides were mapped back to individual proteins and input into the Princeton GO term finder (http://go.princeton.edu/) using all *S. pombe* proteins as background. Process, function, and component analysis were performed.

### CN SLiM prediction

The LxVP motif was defined as [NQDESRTH][YRTDFILV]Lx[VPL][PK] and the PXiXiT motif was defined as P[ACDEFHIKLMNQRSTVWY][IVLF][ACDEFHIKLMNQRSTVWY][IVLF][TSHEDQNKR] as described (Wigington et al., 2020).

### AlphaFold3 structural prediction

Protein structure predictions were generated with the AlphaFold3 server (https://alphafoldserver.com) (Abramson et al., 2024) using protein sequences from Pombase (Carme et al., 2026). The automatic seed setting was used for all predictions. For protein-protein interaction analysis, the top-ranked “model_0” prediction was selected to extract interface information and visualization using the PyMOL molecular graphics system (version 3.0, Schrodinger, LLC). Protein sequence alignments were performed using Clustal Omega (Sievers and Higgins, 2021).

### Statistical analysis

All statistical analyses were performed in Prism 8 (Graphpad software). No data were excluded from the analysis.

## Data availability statement

The data underlying all main and supplemental figures are openly available in Mendeley Data at doi: 10.17632/fchfj9c9fy.1. The mass spectrometry proteomics data have been deposited to the ProteomeXchange Consortium via the PRIDE partner repository with the dataset identifier PXD083328.

## Abbreviations

CR: cytokinetic ring
CN: calcineurin
SLiMs: short linear peptide motifs
IDR: intrinsically disordered region
mNG: mNeonGreen
AIS: autoinhibitory sequence
AF3: AlphaFold3

## Supplemental Table and Figures

**Fig. S1.** mNG-Pxl1 localization in *cdc15* and *pxl1* mutants.

**Fig. S2.** Cdc15 regulation alone cannot explain calcineurin function.

**Fig. S3.** Characterization of Aim21 and Cru1 localization.

**Fig. S4.** Rcn1 is a cytoplasmic protein and dephosphorylated by calcineurin.

**Table S1.** Proteins identified by LC-MS/MS in Ppb1-TurboID.

**Table S2.** Phosphorylation sites identified by LC-MS/MS phosphoproteomic analysis following calcineurin inhibition during cytokinesis.

**Table S3.** Candidate calcineurin substrates localized at the cell division site.

**Table S4.** *S. pombe* strains used in this study.

